# Shoot loss reveals the latent capacity of *Arabidopsis* roots to reconstruct the plant body

**DOI:** 10.64898/2026.09.15.751637

**Authors:** Woo-Taek Jeon, Shawn S. Y. Lee, Changwon Sung, Jung-Min Lee, Ahyeon Cheon, Byungho Kim, Yeonu Lee, Youngsung Joo, Yoo-Sun Noh, Yuree Lee

**Author notes:** Corresponding author: Yuree Lee. These authors contributed equally to this work (Woo-Taek Jeon, Shawn S. Y. Lee). The author responsible for distribution of materials integral to the findings presented in this article in accordance with the policy described in the Instructions for Authors (https://academic.oup.com/plphys/pages/General-Instructions) is: Yuree Lee.

## Abstract

Plant body organization depends on the coordinated development and function of shoots and roots, yet their functional specialization and interdependence remain incompletely understood. Whereas shoot senescence and death, as well as root regeneration from shoot tissues, are well studied, what capacities roots retain in the absence of the shoot remain unclear. To address this question, we tracked Arabidopsis roots after complete shoot excision. Initially, the remaining roots entered a growth-arrested but viable state. Despite broad repression of growth-associated programs, auxin and cytokinin signaling persisted, and the roots retained developmental responsiveness. This arrested state, however, was not developmentally terminal. By approximately 20 days after excision, green structures emerged from lateral root-associated domains. They accumulated chlorophyll and starch, assimilated carbon in response to light, and developed persistent cuticle-covered surfaces. Based on their origin and shoot-like characteristics, we termed them lateral root-derived shoots (LRSs). LRSs arose preferentially in the most shoot-proximal region, where cytokinin signaling progressively accumulated. Excision of this region repositioned LRS formation to the newly established proximal boundary, indicating spatial redefinition of shoot-forming competence. Auxin and cytokinin antagonistically governed the choice between continued lateral root growth and LRS identity, with cytokinin further promoting leaf-like differentiation. LRSs subsequently formed roots and developed into fertile plants under appropriate conditions. These findings reveal that roots retain developmental competence after shoot loss and can reorganize positional and hormonal programs to acquire shoot identity and reconstruct a complete plant body.

## Introduction

The colonization of land exposed plants to contrasting aerial and belowground environments, favoring the differentiation of structures specialized for light and carbon acquisition versus water and mineral uptake (Groff and Kaplan, 1988). In flowering plants, this organization is established early in embryogenesis, when apical–basal patterning specifies the shoot and root meristems, and is subsequently elaborated into distinct organ systems (Lau et al., 2012). Their early specification and persistent functional specialization suggest that shoot and root roles are developmentally stable. Yet the remarkable plasticity of plant development raises a broader question: does this specialization irreversibly constrain organ function, or can established organs deploy developmental capacities beyond their conventional roles?

Despite their distinct developmental origins and functions, shoot and root systems remain physiologically and developmentally interdependent. Carbon fixed in the shoot supports root growth, whereas water and mineral nutrients acquired by roots sustain aerial development, and long-distance signals coordinate these reciprocal activities across the plant body (Puig et al., 2012; Wang and Ruan, 2016; Wheeldon and Bennett, 2021). Auxin and cytokinin are central to this coordination. Shoot-derived auxin contributes to rootward auxin transport, while auxin produced and redistributed locally within roots regulates meristem activity and lateral root development (Roychoudhry and Kepinski, 2022). Root-derived cytokinins participate in long-distance signaling to the shoot, whereas local cytokinin production and signaling shape root growth and differentiation, often through interactions with auxin (Ko and Helariutta, 2017). Shoot and root development are therefore coupled through the integration of organ-local programs with reciprocal metabolic and hormonal signals across the whole plant.

The regenerative capacity of plants, however, shows that established organ identities are not irreversible. Differentiated shoot tissues can acquire root-forming competence after wounding (Shanmukhan et al., 2021; Kareem et al., 2025), while root explants can generate shoots under appropriate culture conditions (Sugimoto et al., 2010). Recent studies indicate that root-to-shoot regeneration can arise through the redirection of pre-existing root developmental programs. Root-derived callus retains features of lateral root development, and lateral root primordia can acquire shoot meristem identity under cytokinin-rich conditions (Atta et al., 2009). These findings demonstrate that cells within established organs can cross conventional shoot–root boundaries. Nevertheless, most experimentally characterized transitions have been induced in excised tissues by exogenous hormones or organogenic culture conditions, leaving unclear whether comparable plasticity can be deployed by an established organ system without direct induction of regeneration.

Natural resprouting provides one context in which belowground organs contribute to recovery after extensive shoot loss. Many perennial species regenerate aerial growth following fire, herbivory, or mechanical damage through the activation of root buds, basal meristems, or other specialized belowground structures (Ott et al., 2019). These systems demonstrate that belowground organs can support whole-plant persistence after extensive shoot destruction, but they often depend on pre-existing buds or organs adapted for storage and resprouting (Clarke et al., 2013). Natural resprouting therefore provides strong evidence for belowground support of shoot recovery but does not establish whether ordinary root tissues lacking preformed shoot structures can generate a new shoot system.

Studies in Arabidopsis have provided a complementary view by showing that roots remain physiologically responsive after shoot removal. Detached roots exhibit enhanced chlorophyll accumulation and chloroplast differentiation when shoot-derived auxin repression is relieved and cytokinin signaling promotes root greening (Kobayashi et al., 2012; Kobayashi et al., 2017). Subsequent work identified type-B ARR-mediated cytokinin signaling and a WIND–B-GATA regulatory pathway as components of shoot removal-induced chloroplast development and photosynthetic gene expression in roots (Kobayashi et al., 2017). Localized non-green callus formation was also observed at the cut surface, spatially distinct from chloroplast development in adjacent root tissues (Kobayashi and Iwase, 2017). Together, these studies established that Arabidopsis roots remain physiologically responsive after shoot removal. However, they left unresolved how the residual root system reorganizes as an integrated developmental system following shoot loss and what trajectory it ultimately follows.

Here, we investigated the early and long-term responses of *Arabidopsis thaliana* roots to complete shoot excision. Shoot-excised roots entered a growth-arrested but viable state while retaining auxin and cytokinin signaling and developmental responsiveness. During prolonged culture, lateral root-associated tissues generated photosynthetically active, cuticle-covered structures that we termed lateral root-derived shoots (LRSs). LRS formation was spatially organized within the residual root system and regulated antagonistically by auxin and cytokinin. LRSs subsequently formed roots and developed into fertile plants, revealing that an established root system can reorganize its developmental programs after shoot loss and reconstruct a complete plant body.

## Results

### Arabidopsis roots remain viable after shoot excision and form lateral root-derived shoots

Natural resprouting illustrates that belowground organs can sustain plant survival and restore aerial growth after severe shoot loss (Fig. 1a and b and S1). However, such recovery often depends on pre-existing buds or specialized belowground structures. To examine how Arabidopsis roots respond to complete shoot loss, we excised the shoots of Arabidopsis thaliana seedlings at 5 days after germination (DAG) and monitored the remaining roots over time (Fig. 1c).

**Figure 1.**
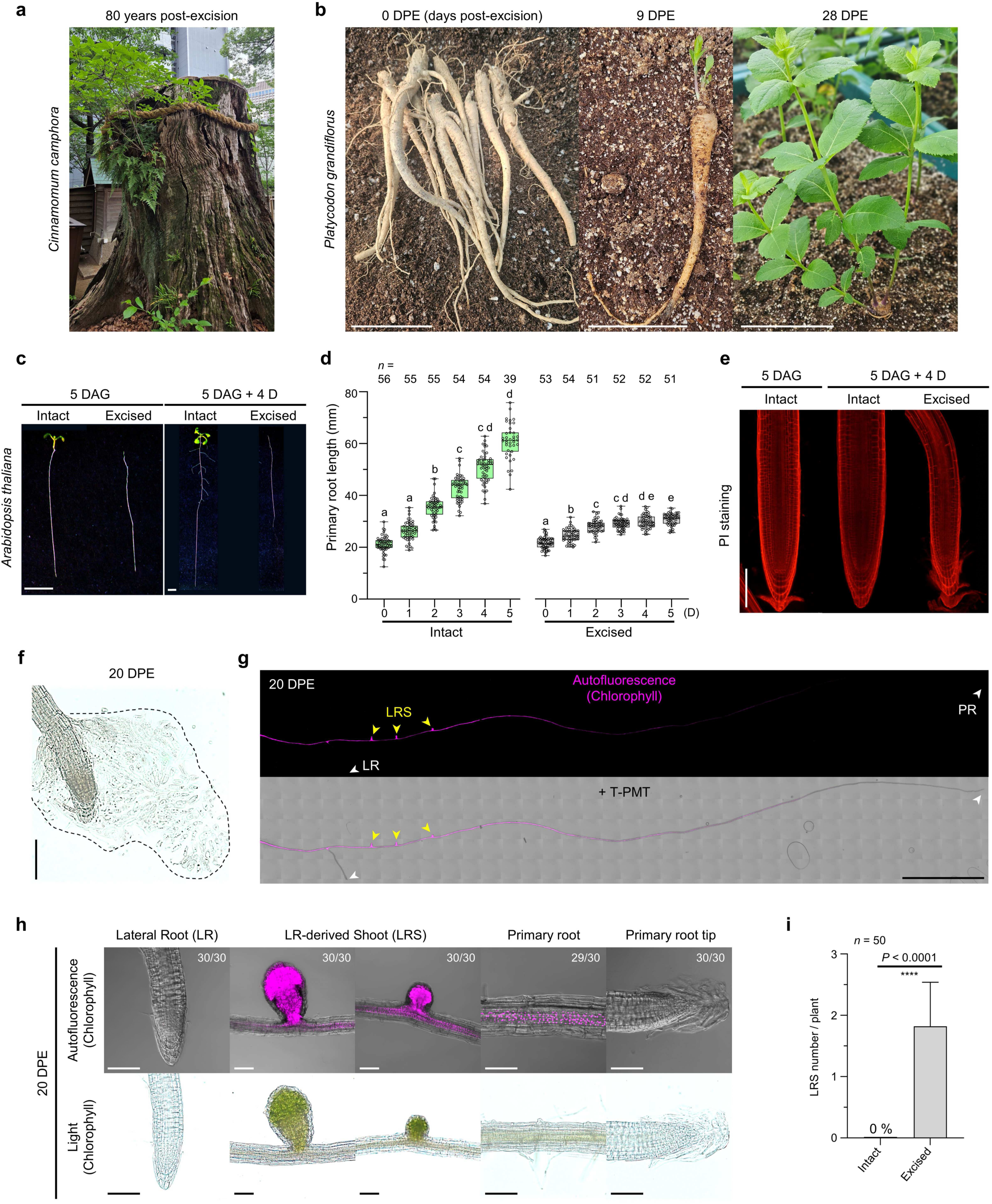
*De novo* shoot regeneration after shoot excision in plants. a) *De novo* shoot regeneration after shoot excision in *Cinnamomum camphora*. The shoot was excised in 1945, and the image was taken in 2025, corresponding to 80 years post excision. b) *De novo* shoot regeneration after shoot excision in *Platycodon grandiflorus*. Plants were maintained after shoot excision in winter 2024 and regrown in spring 2025, during which shoot regeneration was observed. DPE, days post excision. (c) Representative whole-plant images of intact and shoot-excised *Arabidopsis thaliana* plants. Intact indicates untreated plants, whereas excised indicates plants subjected to shoot excision at 5 days after germination (DAG). (d) Quantification of primary root length in the intact and shoot-excised groups. (e) Representative propidium iodide (PI)-stained images of primary root tips at 5 DAG and 4 days after shoot excision in intact and shoot-excised plants (*n* = 30). f) Representative image showing detached root caps from the primary root tip at 20 DPE. g) Overall pattern of lateral root-derived shoot (LRS) formation after shoot excision. Yellow arrowheads indicate LRSs, and white arrowheads indicate lateral roots (LRs) and primary roots (PRs) at 20 DPE. h) Autofluorescence and light microscopy images showing chlorophyll patterning in roots at 20 DPE. i) Quantification of the number of LRSs formed in intact and shoot-excised plants, measured at 25 DAG. Scale bars, 5 cm (b), 5 mm (c), 100 μm (e and h), 200 μm (f), and 500 μm (g). Statistical significance was determined using the Kruskal-Wallis test followed by Dunn’s multiple comparisons test for the intact group in (d), Brown-Forsythe and Welch ANOVA tests followed by Dunnett’s T3 multiple comparisons test for the excised group in (d), and the Mann-Whitney test for (i).

Primary root growth was largely maintained during the first day after excision but declined sharply from 2 days post-excision (DPE). By 3 DPE, root elongation had nearly ceased, with little additional growth thereafter (Fig. 1d). Consistent with this growth arrest, excised roots exhibited reduced meristematic and elongation zones and a progressive decrease in root diameter, whereas roots of intact seedlings continued to grow (Fig. S2a to c).

Despite the pronounced arrest of root growth, the excised roots showed no obvious external signs of tissue deterioration. We therefore examined cellular integrity using propidium iodide (PI) staining at 4 DPE. Shoot-excised roots showed no widespread intracellular PI uptake, indicating that most root cells retained plasma membrane integrity despite the cessation of growth (Fig. 1e). The roots also persisted during extended culture and continued to exhibit root cap cell detachment at 20 DPE, demonstrating that selected developmental activities were maintained long after shoot removal (Fig. 1f). These observations indicate that shoot excision induces a prolonged growth-arrested but viable state rather than rapid root degeneration.

During extended culture, the remaining roots underwent a conspicuous morphological transition. The primary root gradually acquired green pigmentation, and intensely green, rounded structures emerged from lateral root-associated regions (Fig. 1g and h and S2d). Chlorophyll autofluorescence and transmitted-light imaging confirmed strong chlorophyll accumulation in these structures, distinguishing them from the pre-existing primary and lateral roots (Fig. 1g and h). Given their association with lateral root regions and their subsequent acquisition of shoot characteristics, we termed these structures lateral root-derived shoots (LRSs). LRSs were not observed in intact seedlings but formed specifically following shoot excision (Fig. 1i). Together, these findings show that Arabidopsis roots can not only remain viable after shoot excision but also initiate LRS formation at lateral root-associated regions.

### LRSs acquire photosynthetic competence and a persistent cuticle-covered surface

To determine whether LRSs acquire photosynthetic characteristics, we first examined starch accumulation by Lugol’s iodine staining. Because LRSs formed only at a subset of lateral root-associated sites, non-converted lateral roots remained on the same shoot-excised root systems and served as an internal comparison. LRSs exhibited substantially stronger Lugol staining than lateral roots, indicating greater starch accumulation (Fig. 2a and b). Consistent with this result, transmission electron microscopy revealed abundant starch granules within LRS cells (Fig. 2c).

**Figure 2.**
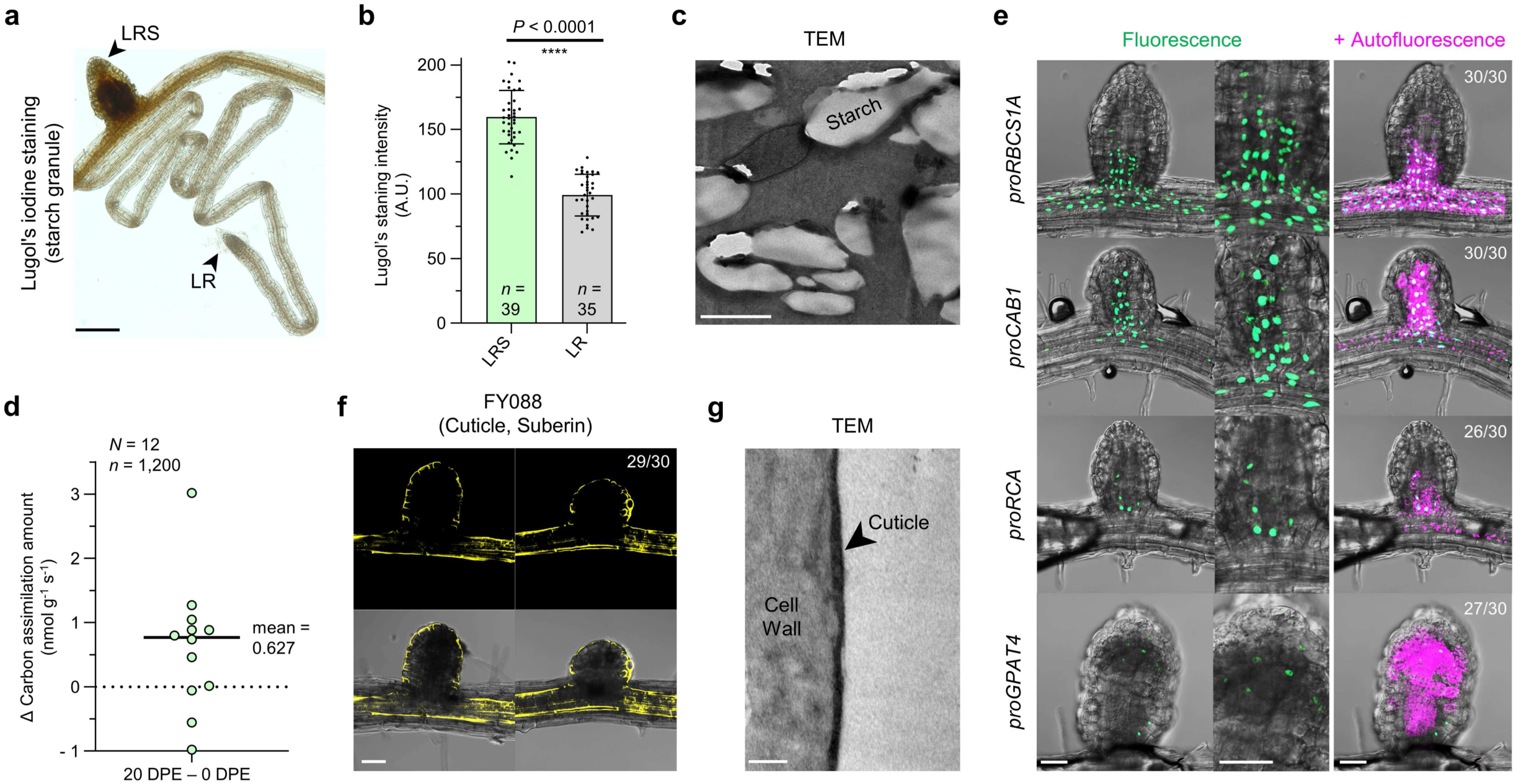
Development of canonical shoot-like structures in LRS. a) Representative images of Lugol’s iodine staining for the detection of starch granules in LRs and LRSs at 20 DPE (*n* > 30). b) Quantification of Lugol’s iodine staining intensity in LRs and LRSs at 20 DPE. c) Representative transmission electron microscopy (TEM) images showing starch granules in LRSs at 20 DPE (*n* = 3). d) Light-induced changes in carbon assimilation. The increase in carbon assimilation at 20 DPE was calculated relative to that at 0 DPE (*n* = 1,200). (e) Representative images showing the activity of photosynthesis- and cuticle-related reporters in LRSs, together with chlorophyll autofluorescence at 20 DPE (*n* = 30). (f) Visualization of cuticle formation in LRSs by Fluorol Yellow 088 (FY088) staining at 20 DPE (*n* = 30). (g) Representative TEM images showing cuticle development in LRSs at 20 DPE. Black arrowheads indicate cuticle (*n* = 3). Scale bars, 200 μm (a), 2 μm (c), 50 μm (e and f), and 400 nm (g). Statistical significance was determined using the Mann–Whitney test for (b).

We next examined whether shoot-excised roots acquired light-dependent carbon assimilation. At 20 DPE, the excised root systems showed a mean increase in carbon assimilation of 0.627 nmol g⁻¹ s⁻¹ relative to 0 DPE, indicating the acquisition of measurable photosynthetic activity during prolonged culture (Fig. 2d). Consistent with this physiological response, reporters driven by the promoters of the photosynthesis-associated genes *RIBULOSE BISPHOSPHATE CARBOXYLASE SMALL CHAIN 1A (RBCS1A)*, *CHLOROPHYLL A/B-BINDING PROTEIN 1 (CAB1)*, and *RUBISCO ACTIVASE (RCA)* were active in LRSs (Fig. 2e). Together with their strong chlorophyll accumulation and starch deposition, these results show that LRSs acquire multiple features associated with photosynthetically active tissues.

We next examined whether LRSs also develop a persistent protective surface. Emerging lateral roots are transiently covered by a cuticle-like surface layer, which is subsequently lost as the root cap detaches during growth (Shin et al., 2025). By contrast, LRSs exhibited a continuous Fluorol Yellow 088-positive surface layer (Fig. 2f). Transmission electron microscopy further revealed an electron-dense layer external to the cell wall, consistent with a cuticle (Fig. 2g). The *GPAT4* promoter, associated with cuticle biosynthesis, was also active in LRSs (Fig. 2e). Thus, unlike mature lateral roots, LRSs maintain a cuticle-covered surface, providing an additional feature characteristic of aerial organs.

### Shoot-excised roots repress growth-associated programs while retaining auxin responsiveness

To characterize the early response of roots to shoot excision, we performed RNA sequencing using primary root tips collected at 1 and 4 DPE, when root growth had largely ceased but LRSs were not yet visible. Total RNA recovered from shoot-excised roots decreased markedly over this period, reaching approximately 10% of the level in intact roots by 4 DPE (Fig. 3a and Table S1). Transcriptome analysis identified 3,597 differentially expressed genes across the examined samples (Fig. 3b). Among these, 1,739 genes that were strongly downregulated after excision were enriched for core biosynthetic and growth-associated processes, including mRNA and rRNA processing, translation, protein folding, and the mitotic cell cycle (Fig. 3c and S3 and Table S2). Many of these genes were highly expressed in intact roots (Table S3), indicating that shoot excision broadly suppresses cellular programs associated with active root growth and biosynthesis.

**Figure 3.**
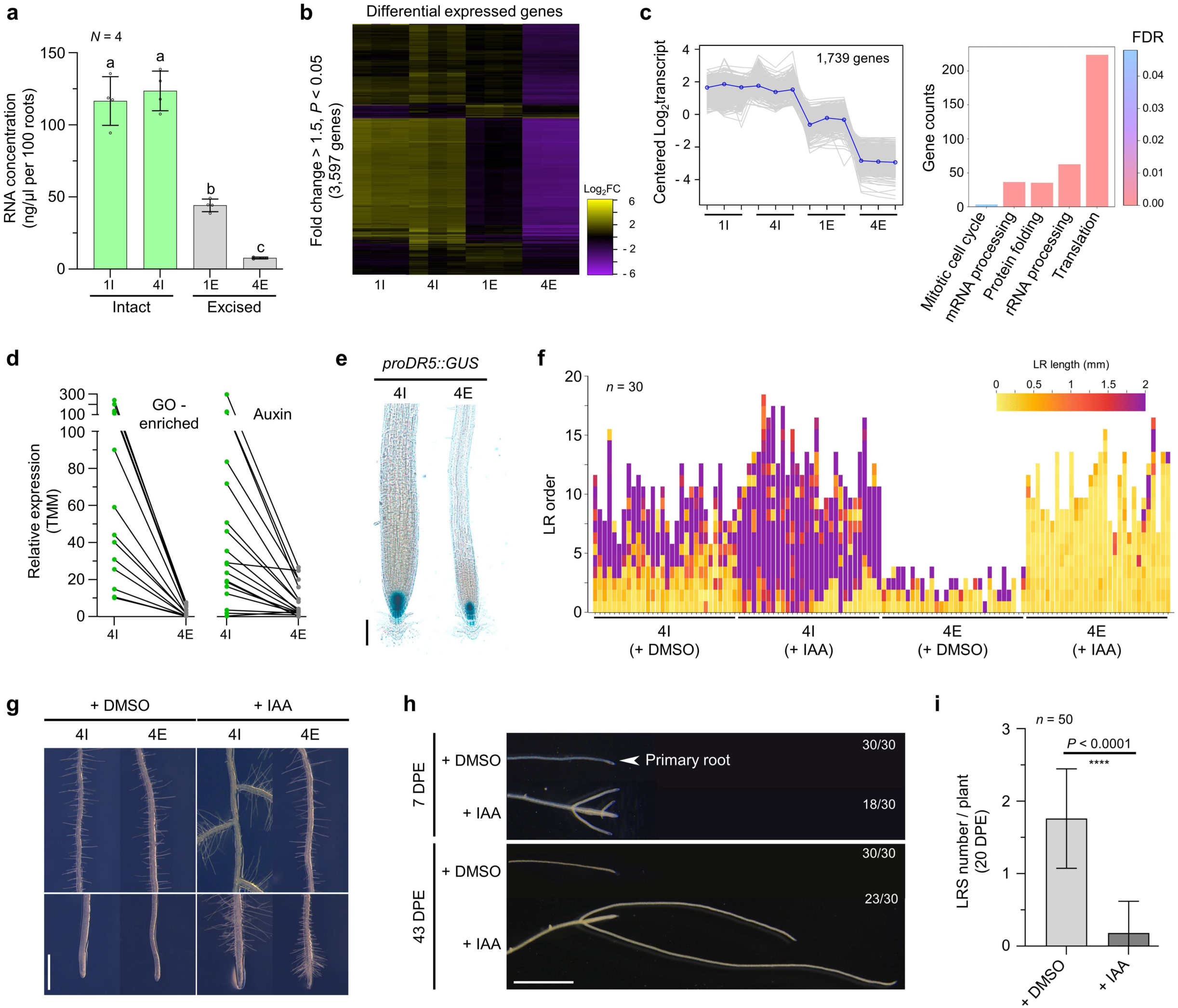
Auxin responsiveness persists after shoot excision but limits LRS development. a) a) RNA concentration extracted from 100 root tips. I and E indicate intact and shoot-excised samples, respectively, and the preceding number indicates days elapsed after 5 days after germination (DAG) (*N* = 4). b) Heatmap of 3,597 differentially expressed genes (DEGs; fold change > 1.50, *P* < 0.05). c) Gene Ontology (GO) analysis of genes downregulated in the shoot-excised group. Bar graphs indicate the number of genes in each category and the corresponding false discovery rate (FDR) for representative GO terms. d) Heatmaps showing the expression patterns of genes associated with the cell cycle, DNA replication, and mRNA processing GO terms, together with auxin-related genes. e) Representative images showing *proDR5::GUS* activity with or without shoot excision (*n* = 30). f) Emerged LR development in response to shoot excision and auxin treatment. Each graph represents an individual plant, and LRs are color-coded according to LR length (*n* = 30). g) Representative images showing root hair patterning in response to shoot excision and auxin treatment. h) LR development in response to auxin treatment after shoot excision at 7 and 43 DPE (*n* = 30). i) Quantification of LRS number in response to auxin treatment after shoot excision. Scale bars, 100 μm (e), 1 mm (g), and 2 mm (h). Statistical significance was determined using ordinary one-way ANOVA for (a) and the Mann-Whitney test for (f).

Notably, auxin-related processes were not overrepresented among genes downregulated after shoot excision (Table S2), suggesting that auxin-associated transcription was relatively preserved despite the broad repression of growth-related programs. Consistent with this interpretation, auxin-related transcripts showed heterogeneous responses and many remained expressed after shoot excision (Fig. 3d and Table S4). The auxin-response reporter *proDR5::GUS* also remained active at the root tip at 4 DPE (Fig. 3e), demonstrating that a localized auxin-response maximum persisted in growth-arrested roots.

We next tested whether the retained auxin signaling was developmentally functional by treating shoot-excised roots with indole-3-acetic acid (IAA). The excised roots remained responsive to IAA across several developmental outputs, including lateral root formation and elongation, root hair development, primary root growth, and root cap detachment (Fig. 3f and g and Fig. S4). During prolonged culture, IAA did not restore sustained primary root elongation but promoted continued lateral root growth (Fig. 3h). In contrast, exogenous IAA strongly suppressed LRS formation (Fig. 3i). Thus, although shoot excision broadly represses growth-associated cellular programs, the remaining roots retain functional auxin-response pathways. Auxin preferentially supports continued lateral root development while antagonizing the transition toward LRS formation.

### Cytokinin signaling promotes LRS formation within a dynamically positioned shoot-proximal domain

Cytokinin is a central regulator of root–shoot coordination and *de novo* shoot organogenesis (Ikeuchi et al., 2016). Given that auxin promoted continued lateral root development while suppressing LRS formation, we next examined whether cytokinin contributes to LRS development. Cytokinin-related transcripts remained detectable after shoot excision, although many showed reduced expression (Fig. 4a and Table S4). The cytokinin-response reporter *proTCSn::nTdTomato* also remained active in the primary root tip at 4 DPE (Fig. 4b). These observations indicate that cytokinin signaling persists in growth-arrested roots following shoot removal.

**Figure 4.**
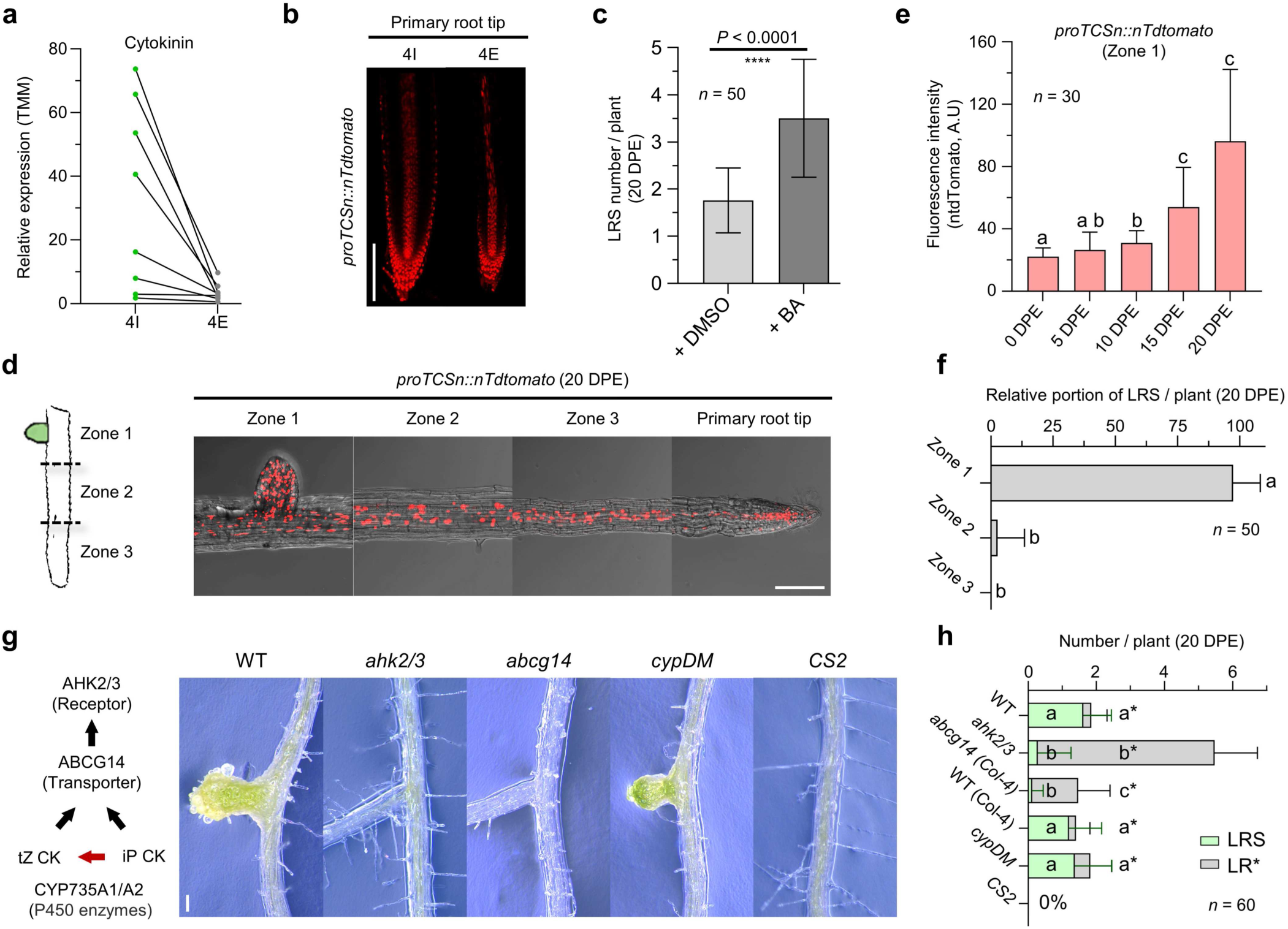
Cytokinin sink formation is required for LRS initiation. a) Heatmaps showing the expression patterns of cytokinin-associated genes after shoot excision. b) Representative images showing *proTCSn::nTdtomato* activity with or without shoot excision (*n* = 30). c) Quantification of LRS number in response to exogenous cytokinin treatment with 6-benzylaminopurine (BA). (d) Representative images showing *proTCSn::nTdtomato* activity at 20 DPE (*n* = 30). Zone 1, Zone 2, and Zone 3 correspond to the upper, middle, and lower thirds of the primary root, respectively. e) Quantification of *proTCSn::nTdtomato* signal intensity in Zone 1 over time after shoot excision (*n* = 30). f) Quantification of the proportion of LRSs formed in each zone within individual plants (*n* = 50). g) Representative images showing LRS development in cytokinin- and lateral root-related mutants (*n* = 30). *CS2* indicates *proCASP1::shy2-2* and *cypDM* indicates *cyp735a1-2 cyp735a2-2* double mutant. h) Quantification of LR and LRS numbers in cytokinin- and lateral root-related mutants (*n* = 60). Scale bars, 100 μm (b and e). Statistical significance was determined using the Kruskal-Wallis test followed by Dunn’s multiple comparisons test for (e), (f), and (h).

To test its functional contribution, we treated shoot-excised roots with the synthetic cytokinin 6-benzyladenine (BA). BA treatment significantly increased the number of LRSs, demonstrating that cytokinin promotes LRS formation (Fig. 4c). We then examined the spatial distribution of cytokinin signaling during prolonged culture. The primary root was divided into three regions, designated Zone 1, Zone 2, and Zone 3 from the shoot-proximal end toward the root apex (Fig. 4d). The *proTCSn::nTdTomato* signal persisted throughout the excised root and progressively increased in Zone 1, particularly at later time points (Fig. 4d and e). Consistent with this spatial pattern, LRSs formed predominantly within Zone 1 (Fig. 4f), indicating that LRS formation is associated with a cytokinin-responsive shoot-proximal domain.

We next tested whether this domain was tied to a fixed position within the original root axis or defined relative to the newly established proximal boundary. Removal of Zone 1 reduced overall LRS formation, but the LRSs that subsequently formed arose preferentially within the newly proximal Zone 2 region (Fig. S5a and b). Thus, shoot-forming competence is not restricted to a permanently specified root segment but can be repositioned toward the most shoot-proximal region remaining after excision.

Genetic analyses further supported the requirement for cytokinin perception and transport in LRS formation. LRSs were not detected in the cytokinin receptor mutant *ahk2 ahk3* or in the cytokinin transporter mutant *abcg14* under our conditions (Fig. 4g and h). By contrast, the *cyp735a1 cyp735a2* double mutant, which is impaired in CYP735A-dependent production of trans-zeatin-type cytokinins, formed LRSs at levels comparable to those of wild type (Fig. 4g and h). This result indicates that CYP735A-dependent trans-zeatin production is not essential for LRS formation and suggest that other cytokinin forms can support this process. This interpretation is consistent with evidence that ABCG14 contributes to the transport of both iP- and trans-zeatin-type cytokinins (Zhao et al., 2023). Finally, LRSs were absent from the lateral root-defective *CS2* line (*proCASP1::shy2-2*), in which lateral root formation was strongly reduced (Fig. 4g and h). This result supports the conclusion that LRS formation depends on lateral root development or lateral root-associated developmental domains.

Together, these findings show that cytokinin promotes LRS initiation within a dynamically positioned shoot-proximal domain and that this process depends on cytokinin perception and transport as well as lateral root-associated developmental competence.

### Cytokinin promotes leaf-like differentiation of LRSs and regeneration into fertile plants

To determine whether cytokinin influences LRS development beyond its initial formation, we examined the morphology of LRSs following BA treatment. BA-treated LRSs were markedly larger than control LRSs and developed pronounced leaf-like structures (Fig. 5a). Despite this enhanced growth and differentiation, BA did not broaden the spatial domain of LRS formation. Under both control and BA-treated conditions, LRSs remained predominantly restricted to Zone 1 (Fig. 5b). Thus, cytokinin promotes the development of LRSs within the pre-established shoot-proximal domain rather than extending LRS-forming competence along the primary root.

**Figure 5.**
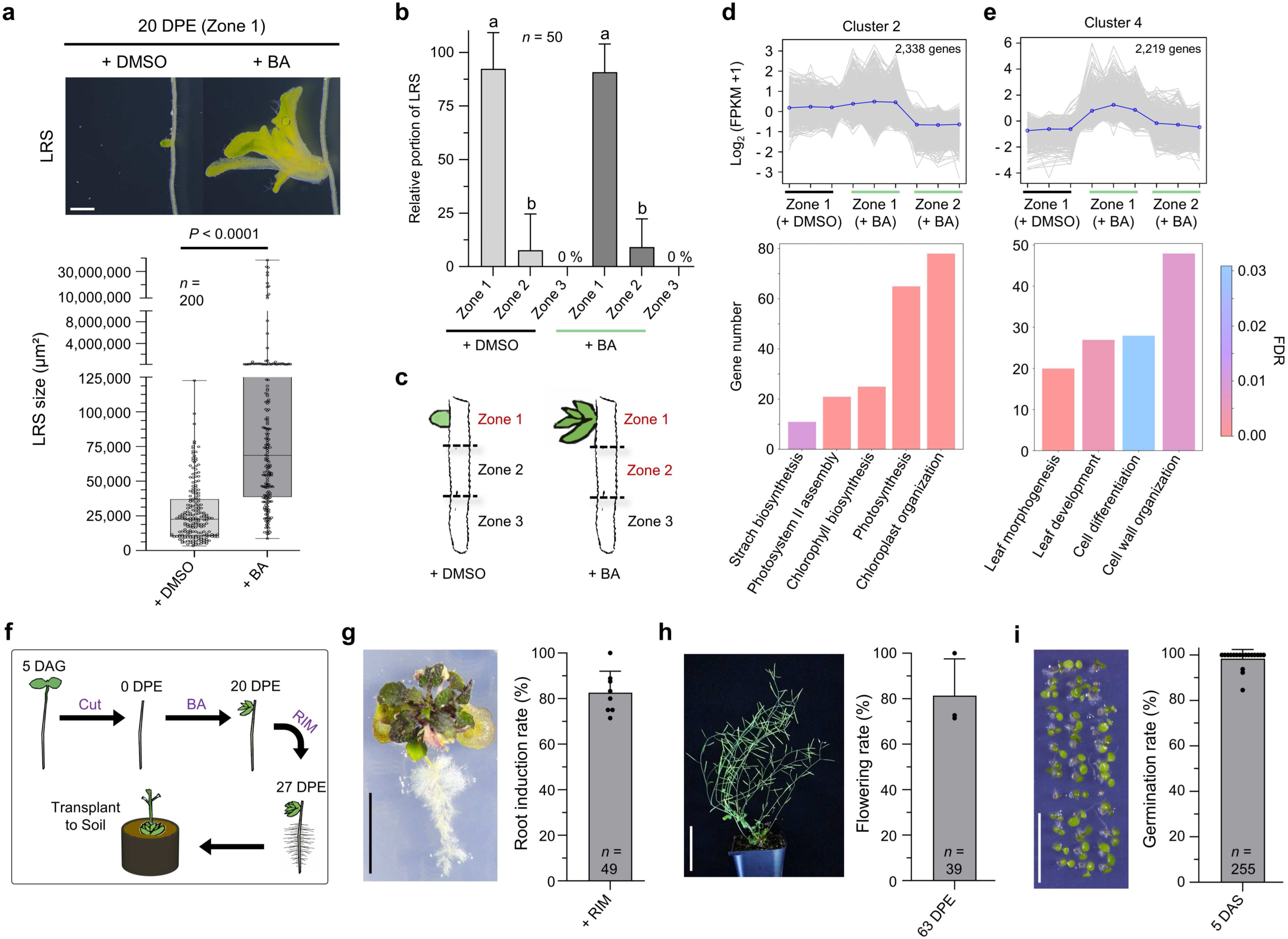
Cytokinin drives LRS maturation and enables whole-plant regeneration. a) Representative images and quantification of LRS development in response to exogenous cytokinin treatment with 6-benzylaminopurine (BA) (*n* = 200). b) Proportion of LRSs formed in each zone within individual plants in response to BA treatment. c) Schematic diagram of the RNA-seq experimental design. Samples indicated in red were subjected to sequencing. d) Gene Ontology (GO) analysis of genes upregulated in Zone 1 independently of BA treatment among differentially expressed genes (DEGs; P < 0.05, fold change > 1.5). Bar graphs indicate the number of genes in each category and the corresponding false discovery rate (FDR) for representative GO terms. e) GO analysis of genes specifically upregulated in Zone 1 after BA treatment among DEGs. f) Schematic diagram of the experimental workflow for whole-plant regeneration from LRSs. g) Representative images showing root induction after 7 days of incubation on root induction medium (RIM), together with quantification of the root induction rate. h) Representative whole-plant images after transplantation to soil at 63 DPE, together with quantification of the flowering rate. i) Representative images of next-generation seedlings obtained from regenerated plants, together with quantification of the germination rate at 5 days after sowing (DAS). Scale bars, 500 μm (a), 1 cm (g and i), and 5 cm (h). Statistical significance was determined using the Mann-Whitney test for (a) and the Kruskal-Wallis test followed by Dunn’s multiple comparisons test for (b).

To examine the transcriptional basis of this spatial restriction, we performed RNA sequencing using Zone 1 from control roots and Zones 1 and 2 from BA-treated roots (Fig. 5c and Table S5). A cluster of 2,338 genes showed higher expression in Zone 1 under both control and BA-treated conditions than in BA-treated Zone 2 (Fig. 5d). Gene Ontology enrichment analysis showed that this cluster was associated with starch biosynthesis, chlorophyll biosynthesis, photosynthesis, photosystem assembly, and chloroplast organization (Fig. 4d and Table S6). These results indicate that photosynthetic and chloroplast-related programs are intrinsic features of the shoot-proximal LRS-forming domain and remain spatially restricted even under elevated cytokinin conditions.

We next identified transcriptional changes associated with the enhanced leaf-like morphology of BA-treated LRSs. A separate cluster of 2,219 genes was preferentially expressed in BA-treated Zone 1 relative to both control Zone 1 and BA-treated Zone 2 (Fig. 5e). This cluster was enriched for genes associated with leaf morphogenesis, leaf development, cell differentiation, and cell wall organization (Fig. 5e and Table S7). Consistently, genes related to leaf development, epidermal identity, and cuticle formation were preferentially expressed in BA-treated Zone 1 (Fig. S5c). These findings indicate that cytokinin acts within the shoot-proximal domain to promote the differentiation of LRSs toward a leaf-like developmental state.

Finally, we tested whether LRS-derived shoots could regenerate complete plants (Fig. 5f). Following BA-mediated shoot development, LRS-derived shoots were transferred to root induction medium, where most formed roots (Fig. 5g and Fig. S5d to f). The rooted plants subsequently grew in soil and produced flowers (Fig. 5h). Seeds obtained from these regenerated plants germinated and developed into the next generation (Fig. 5i). Thus, LRS-derived shoots retain the developmental capacity to regenerate fertile plants, demonstrating that roots can reconstruct a complete plant body after shoot loss.

## Discussion

Complete shoot loss would be expected to severely compromise root development because roots normally depend on the shoot for photosynthetically derived carbon and long-distance developmental signals. Our findings instead reveal that the remaining root system retains substantial developmental capacity. Shoot-excised roots entered a prolonged growth-arrested but viable state, from which lateral root-associated tissues subsequently generated LRSs. LRSs acquired photosynthetic competence and a persistent cuticle-covered surface, and their formation was spatially restricted within the residual root system and differentially regulated by auxin and cytokinin. Cytokinin treatment further promoted leaf-like differentiation, after which LRS-derived shoots could be guided through root induction to regenerate fertile plants. These findings indicate that shoot loss does not immediately exhaust root developmental potential but exposes a staged process in which retained root competence can be redirected toward aerial organ formation and, under appropriate hormonal and culture conditions, complete plant regeneration (Fig. 6).

**Figure 6.**
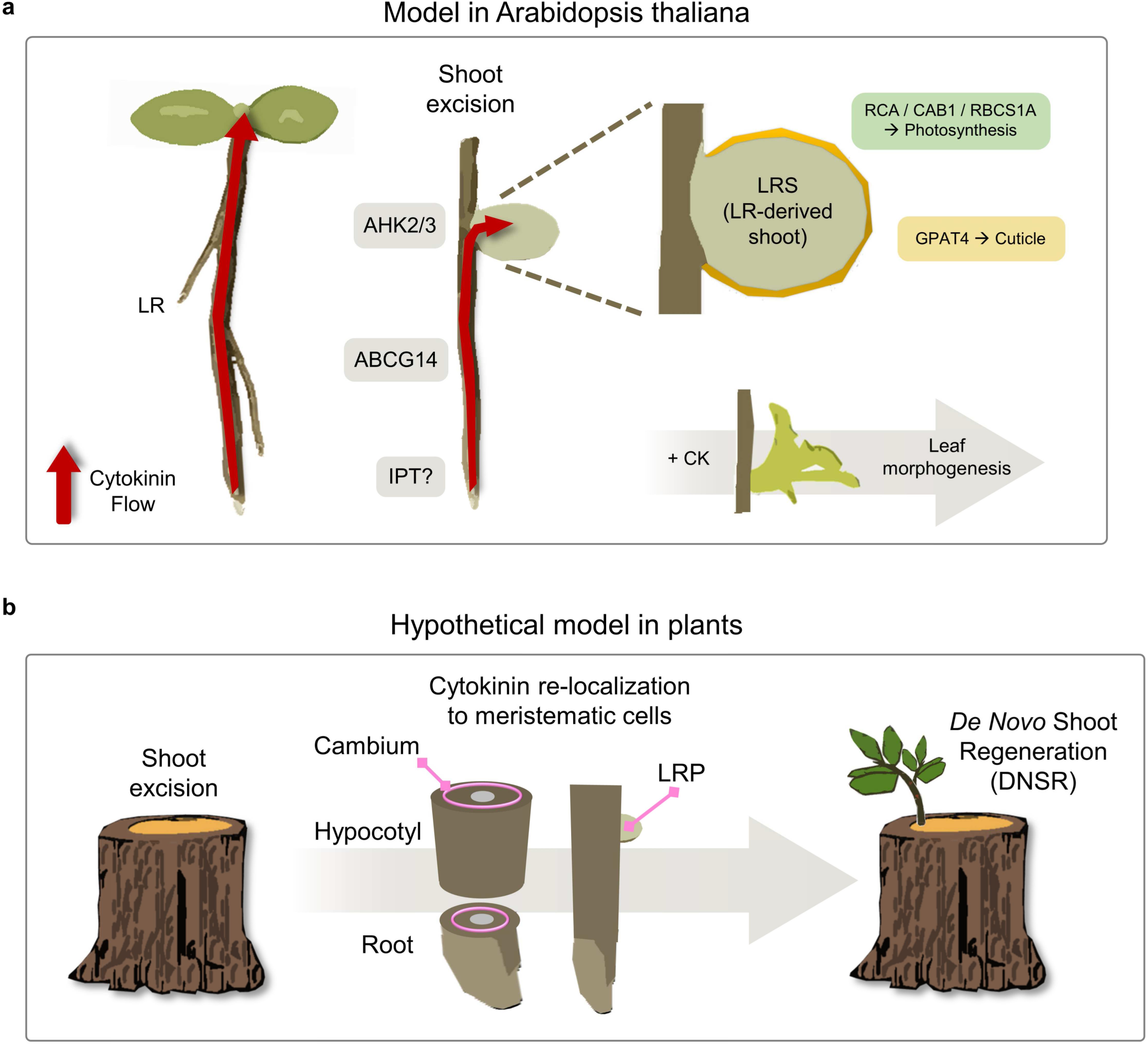
Model for *de novo* shoot regeneration in plants. a) Schematic model of LRS-mediated *de novo* shoot regeneration in Arabidopsis. Under normal developmental conditions, root-derived cytokinin is transported toward the shoot organ, which acts as a cytokinin sink. After shoot excision, cytokinin is redirected and accumulates at lateral root primordia, promoting lateral root-derived shoot (LRS) development. Cytokinin transport requires ABCG14, and cytokinin perception requires AHK2 and AHK3. During LRS development, photosynthesis-related genes are expressed in inner cells, leading to chlorophyll accumulation, whereas cuticle formation occurs in the outermost epidermal layer, together conferring shoot-like properties. Exogenous cytokinin treatment further promotes complete leaf morphogenesis. b) Hypothetical model of *de novo* shoot regeneration in plants. After shoot excision, cytokinin sinks may be relocated to meristematic tissues, such as the cambium or lateral root primordia, thereby promoting shoot development.

A central feature of the response to shoot loss was selective preservation rather than uniform developmental shutdown. Shoot-excised roots strongly repressed the biosynthetic, cell-cycle, and elongation programs required for sustained primary root growth, while retaining localized hormonal responses and selected developmental functions. Exogenous IAA supported lateral root development but failed to restore sustained primary root elongation, indicating that growth capacity was differentially retained across the root system. Particularly notable was the preservation of lateral root primordia or associated organogenic domains, which later served as the sites of LRS initiation. Shoot loss therefore appears to suppress continued expansion of the primary axis while sparing specific meristematic domains with the potential to generate new organs. This selective response raises a fundamental question: what molecular mechanisms distinguish developmental competence that should be preserved from growth programs that can be curtailed? The shoot-excised root system may therefore provide a useful model for identifying the transcriptional and epigenetic mechanisms that selectively preserve organogenic competence during growth arrest.

The developmental significance of this selective preservation became evident when organogenesis resumed, not through renewed elongation of the primary root axis, but through the formation of LRSs. At LRS-forming sites, lateral root-associated primordia diverged from the conventional trajectory of root cap maturation and sustained axial elongation. Instead, they expanded into enlarged structures and progressively acquired shoot-associated characteristics. This divergence provides an important connection to in vitro regeneration, in which callus induction reactivates lateral root or root meristem programs that subsequently support shoot formation under cytokinin-rich conditions (Sugimoto et al., 2010). The lateral root-like state of callus may therefore reflect an inherent capacity of root organogenic programs to be redirected toward shoot development. However, the cellular heterogeneity and spatial disorganization of callus make it difficult to reconstruct the trajectory of an individual organogenic unit. Because LRSs arise at discrete lateral root-associated sites within an intact root axis, they provide a primordium-resolved system for following the divergence of root organogenesis toward shoot formation.

Current regeneration frameworks suggest that this transition may involve molecularly distinct phases. Lateral root initiation and auxin-induced callus formation engage root meristem-associated regulators, including LBD and PLETHORA modules, whereas cytokinin-dependent shoot regeneration recruits type-B ARRs and shoot meristem regulators such as WUSCHEL, CUP-SHAPED COTYLEDON, and SHOOT MERISTEMLESS (Fan et al., 2012; Kareem et al., 2015; Meng et al., 2017). Chromatin accessibility and Polycomb-associated repression further influence the acquisition and expression of regenerative competence (He et al., 2012; Wu et al., 2022). The LRS system offers an opportunity to determine how these regulatory layers are temporally coordinated—from the preservation of lateral root-associated competence during growth arrest to the activation of shoot-organogenic programs. Such analysis should clarify whether LRS formation redeploys established regeneration circuits or reveals a more direct route by which a lateral root-associated organogenic domain acquires shoot identity.

This developmental redirection was not deployed uniformly across the residual root system but was strongly constrained by position. Cytokinin signaling progressively increased in the shoot-proximal region, suggesting that the establishment of a local cytokinin-responsive domain contributes to the spatial bias of LRS formation. Cytokinin-mediated source–sink regulation provides a possible physiological context for this pattern, as cytokinin allocation can be redirected when dominant shoot sinks are lost or weakened (Ko and Helariutta, 2017; Walker et al., 2023; Jeon et al., 2026). The requirement for ABCG14 is consistent with a contribution of cytokinin transport or redistribution to the emerging proximal domain. However, exogenous BA enhanced LRS initiation and differentiation without extending the competent region along the primary root, indicating that cytokinin alone is insufficient to specify LRS position. Moreover, removal of the original proximal region repositioned LRS formation toward the newly established proximal boundary. Together, these observations support a model in which shoot loss redefines positional competence relative to the residual root axis, while cytokinin reinforces this domain and promotes LRS development. During embryogenesis, apical–basal polarity is established through spatially organized auxin transport and transcriptional networks that distinguish shoot and root poles (Lau et al., 2012). Whether components of these polarity mechanisms are redeployed to establish a new shoot-forming domain after shoot loss remains an important question. Because our transcriptomic analysis captured the established proximal state rather than its initial specification, analysis of the earlier response may reveal the transcriptional and epigenetic modules that redefine axial information within an existing root system.

Taken together, our findings suggest that plant body reconstruction after organ loss may rely substantially on the reassignment of pre-existing meristematic competence. In Arabidopsis roots, lateral root-associated domains provide such a developmental substrate, enabling a root organogenic program to be redirected toward aerial development and, under appropriate culture conditions, whole-plant regeneration. More broadly, loss of a dominant shoot may alter hormonal and source–sink relationships in ways that recruit alternative meristematic compartments, such as root primordia, basal tissues, or cambial domains, to initiate new shoot-forming programs (Fig. 6). Whether this principle contributes to natural resprouting or *de novo* shoot regeneration in other species remains to be tested. The LRS system therefore offers a tractable model for determining how plants preserve, reassign, and execute developmental potential following severe organ loss.

## Materials and Methods

### Plant materials and growth conditions

*Arabidopsis thaliana* lines used in this study were derived from the Columbia-0 (Col-0) ecotype, except for *abcg14* and its corresponding wild-type control, which were in the Columbia-4 (Col-4) background. Detailed information on the *Arabidopsis* mutant and transgenic lines used in this study has been provided in previous publications: *proRBCS1A::nlsGFP-GUS*, *proCAB1::nlsGFP-GUS*, *proRCA::nlsGFP-GUS* (Wen et al., 2025), *proGPAT4::nlsGFP-GUS* (Lee et al., 2025), *proDR5::GUS*, *proUBQ10::4xYFP* (Shin et al., 2025), *proTCSn::ntdTomato* (Smet et al., 2019), *ahk2-2 ahk3-3* (Riefler et al., 2006), *cyp735a1-2 cyp735a2-2* (Kiba et al., 2013), *abcg14* (Ko et al., 2014), *proCASP1::shy2-2* (Vermeer et al., 2014). Seedlings were grown on half-strength Murashige and Skoog (MS) medium supplemented with 1% (w/v) sucrose and solidified with agar. Seeds were surface sterilized using a vapor-phase method and stratified at 4°C for 2 days before sowing. Plants were grown under long-day conditions consisting of 16 h light at 22°C and 8 h darkness at 18°C, with a light intensity of 70 μmol m^−2^ s^−1^.

Wild *Platycodon grandiflorus* plants were collected from natural populations and subsequently cultivated in a greenhouse maintained at 20°C–22°C under natural light conditions. *Cinnamomum camphora* (Kusunoki no Shinboku) growing at Ikuta Shrine, Kobe, Japan, was observed and photographed using a Samsung Galaxy S25 (Samsung Electronics, Suwon, South Korea).

### Shoot excision assay

At 5 DAG, the entire shoot, including the hypocotyl, shoot apical meristem, and leaves, was removed with a single clean cut using sterile scissors in a laminar flow hood. For excision treatments that included Zone 1, the upper one-third of the primary root was additionally removed. Immediately after excision, the culture plates were resealed to prevent desiccation of the medium and returned to the same growth conditions.

### Pharmacological assays

For auxin and cytokinin treatments, seedlings were initially germinated and grown on hormone-free half-strength MS medium. At the indicated time points, seedlings were transferred to medium supplemented with indole-3-acetic acid (IAA) or 6-benzyladenine (BA). For experiments involving shoot excision, seedlings were transferred to the hormone-containing medium immediately after excision. IAA and BA were dissolved in dimethyl sulfoxide (DMSO) and added to the medium at a final concentration of 4 μM.

### Phenotypic assessment

Images of *Cinnamomum camphora* and *Platycodon grandiflorus*, as well as germinating Arabidopsis seedlings, whole plants, and reproductive architecture, were captured using a Samsung Galaxy S25 smartphone (Samsung Electronics, Suwon, South Korea). Inflorescences and fertile and sterile fruits were counted individually.

For measurement of total root length, plants were photographed while positioned on the growth medium using a KCS3-50 camera (Korea Lab Tech, Gyeonggi-do, South Korea). Root length was subsequently quantified from the images using ImageJ (Schneider et al., 2012).

### Microscopy and visualization

Bright-field images of root cap detachment, root MZ, root EZ, root diameter, root greening, LRS number, LR number, Lugol-stained roots, and *DR5-GUS*-expressing roots were acquired using an Axio Observer 5 microscope (Zeiss, Jena, Germany). Root hair patterns and root growth patterns were examined using an M205 FA stereo microscope (Leica, Wetzlar, Germany), which was also used to acquire LRS images for size measurement. Confocal imaging was performed using an LSM 900 confocal microscope (Zeiss, Jena, Germany) with the following excitation wavelengths and detection ranges: 561 nm excitation and 580–680 nm detection for propidium iodide (PI), 488 nm excitation and 500–545 nm detection for GFP, 561 nm excitation and 580–630 nm detection for tandemTomato (tdT), and 488 nm excitation and 650–750 nm detection for chlorophyll autofluorescence.

Plants harboring the *DR5::GUS* reporter construct were harvested and briefly rinsed in GUS staining buffer containing 50 mM sodium phosphate buffer (pH 7.2), 0.2% (v/v) Triton X-100, 2 mM potassium ferrocyanide, and 2 mM potassium ferricyanide. The buffer was then replaced with fresh GUS staining solution supplemented with 10 mM X-Gluc (Duchefa Biochemie), and samples were incubated for 2 h at 37°C in the dark. The enzymatic reaction was terminated by immersing the samples in 100% ethanol for 1 h, which also facilitated tissue clearing. Plants were depigmented overnight in 100% ethanol to completely remove chlorophyll and subsequently stained with Lugol’s iodine solution containing 6 mM iodine, 43 mM KI, and 0.2 N HCl for 1 min to visualize starch granules. The roots were then gently rinsed with distilled water to remove excess background staining. Plasma membrane integrity was assessed by incubating samples in 10 μM propidium iodide (PI) for 2 min, followed by rinsing with water twice. Cuticle staining with Fluorol Yellow 088 was performed using a modified version of the method described by (Jeon et al., 2026). Root samples were immersed in 0.01% (w/v) Fluorol Yellow 088 dissolved in methanol and incubated for 7 days at room temperature in darkness. Samples were then stained with 0.5% (w/v) aqueous aniline blue for 1 hour to suppress nonspecific background fluorescence, washed thoroughly with water, and examined by fluorescence microscopy.

### Photosynthesis Capacity Quantification

A portable infrared gas analyzer (GFS-3000, Walz, Jena, Germany) was used to measure the root carbon assimilation rate. For each biological replicate, 100 root segments were collected and their fresh weights were recorded for normalization. Measurements were made at 600 µmol s^−1^ of flow, 440 ppm of CO2 concentration, and a cuvette temperature of 22 °C. The carbon assimilation rate for each replicate was calculated by comparing the gas exchange at PAR = 0 µmol m^-2^ s^-1^ and $PAR = 100 µmol m^-2^ s^-1^. The measurement at PAR = 0 was used to estimate the net respiration rate, while the value at PAR = 100 estimated the combined rate of respiration and photosynthesis. To quantify the specific gain in photosynthetic capacity during regeneration, the mean carbon assimilation of the control group (0 DPE) was subtracted from the values of each replicate in the treatment group (20 DPE). The final increased photosynthesis amount was expressed in nmol g^-1^ s^-1^.

### Transmission electron microscopy

For transmission electron microscopy (TEM), tissue specimens were immersed overnight at 4°C in a fixative containing 2% paraformaldehyde and 2% glutaraldehyde prepared in 50 mM potassium phosphate buffer (pH 7.0). The samples were then post-fixed with 1% (w/v) osmium tetroxide for 2 h at 4°C. After three rinses with distilled water, the specimens were dehydrated through an ethanol series of 30%, 50%, 70%, 80%, and 90% for 30 min at each concentration, followed by three 30-min incubations in 100% ethanol. The dehydrated tissues were infiltrated with Spurr resin (Electron Microscopy Sciences, Pennsylvania, USA) and cured at 70°C for 48 h. Ultrathin sections, approximately 50 nm in thickness, were collected and observed using a Tecnai G2 Spirit transmission electron microscope (FEI, Oregon, USA) operated at 120 kV under high-vacuum conditions.

### Whole-plant regeneration and physiological assessment of LRS-derived plants

To induce lateral root–derived shoots (LRSs), shoots were mechanically excised from 5 DAG. Immediately after excision, the remaining root tissues were transferred to half-strength MS medium supplemented with 4 μM BA and cultured for 20 days to induce leaf-like morphogenesis from LRSs. For root induction, intact explants with regenerated shoots were placed onto the root induction medium (RIM). RIM used in this study contains Gamborg’s B5 medium with minimal organics (Duchefa Biochemie), 3% sucrose buffered to pH 5.7 with 0.05% MES, and 0.8% phytoagar supplemented with 158 mg/l indole-3-acetic acid (IAA). The base of each regenerated shoot was gently embedded into a shallow hole prepared with forceps, keeping the shoot apex exposed above the medium surface. Successfully rooted plantlets were transferred to soil, and their subsequent growth and development were monitored. The numbers of inflorescences and fertile and sterile seeds were determined at 63 DPE, corresponding to 68 DAG. Seeds harvested from the regenerated plants were sown on half-strength MS medium, and germination rates were assessed 5 days after sowing.

### Transcriptomic analysis

Samples were obtained from three independent experimental replicates. For root-tip transcriptome analysis, 12 samples were prepared, with each sample comprising primary root tips collected by excising approximately 0.5 cm from the root apex. For LRS transcriptome analysis, 9 samples were prepared, each containing material collected from roots at the indicated regions. Total RNA was isolated using the RNeasy Plant Mini Kit (Qiagen, Hilden, Germany) according to the manufacturer’s instructions. RNA quality was assessed before library preparation, and samples with an RNA integrity number (RIN) >9 were used for RNA sequencing. Stranded mRNA libraries were generated using the TruSeq Stranded mRNA Library Prep Kit (Illumina, California, USA) following the manufacturer’s protocol and sequenced on an Illumina HiSeq 2500 platform (Illumina, California, USA).

RNA-sequencing data were processed using the VizR platform with the following parameters and analysis settings (Jeon et al., 2026). TruSeq universal and indexed adapter sequences were removed using Trimmomatic v0.39 (Bolger et al., 2014), and the resulting reads were quality-filtered using PRINSEQ v0.20.4 (Schmieder and Edwards, 2011). Filtered reads were aligned to the Arabidopsis TAIR10.1 reference genome (GCF_000001735.4) using HISAT2 v2.2.1 (Kim et al., 2019). Gene abundances were quantified using StringTie v1.3.4 with the corresponding TAIR10 GFF3 annotation file, and expression levels were summarized as transcripts per million (TPM) (Pertea et al., 2015). For conventional between-sample normalization, trimmed mean of M-values (TMM) normalization was performed using edgeR v3.19 in R (Grabherr et al., 2011).

An additional normalization procedure was applied to the root-tip RNA-sequencing dataset because total RNA yields were markedly reduced following shoot excision. Consequently, conventional TMM normalization and normalization incorporating the experimentally measured RNA amount produced substantially different expression patterns (Fig. S6a). To determine which normalization approach more appropriately represented the biological response, the expression pattern of *UBQ10* was used as an internal reference and compared with its corresponding expression pattern in the RNA-sequencing dataset (Fig. S6, b to d). This comparison supported normalization based on RNA amount rather than TMM normalization for the root-tip dataset. Accordingly, RNA amount-normalized expression values were used for subsequent analyses of the root-tip transcriptome (Table S1). In contrast, no statistically significant difference in RNA yield was detected among the LRS samples, which were collected under comparable shoot-excision conditions; therefore, conventional TMM normalization was applied to the LRS dataset (Fig. S6e and Table S5)

Differentially expressed genes (DEGs) were identified using thresholds of fold change >1.5 and P < 0.05. Expression patterns of selected genes were visualized as row-wise Z-score heatmaps using the pheatmap package v1.0.12 in R (Kolde and Kolde, 2015). Gene clustering was performed using the Ptree method implemented in Trinity (Grabherr et al., 2011). Clusters showing similar expression trends were subsequently combined, and the resulting gene sets were subjected to Gene Ontology (GO) enrichment analysis using DAVID (Sherman et al., 2022).

### Quantification and statistical analysis

Staining intensity and stained area were quantified using Fiji (Schindelin et al., 2012). Details of biological replicates and the statistical tests applied are described in the corresponding figure legends. Statistical analyses were conducted using GraphPad Prism version 10.5.0.

## Acknowledgements

We thank Professor June M. Kwak for the *proRBCS1A::nlsGFP-GUS*, *proCAB1::nlsGFP-GUS*, and *proRCA::nlsGFP-GUS* lines, Professor Niko Geldner for the *proCASP1::shy2-2* line, Professor Bert De Rybel for the *proTCSn::ntdTomato* line, Professor Hyunwoo Cho for providing the *ahk2-2 ahk3-3* mutant and Dr. Donghwi Ko for the *cyp735a1-2 cyp735a2-2* and *abcg14* mutants. Yuree Lee was supported by the Suh Kyungbae Foundation (SUHF-19010003), Mid-Career Bridging Program from Seoul National University and the National Research Foundation of Korea (NRF) grant funded by the Korea government (No.RS-2021-NR60084 and RS-2026-25491168). YS Noh was supported by grants from the NRF grant (No.RS-2021-NR060084). W-T Jeon was supported by the Presidential Science Scholarship and NRF grant (No. RS-2026-25553773). W-T Jeon and J-M Lee were supported by the Stadelmann–Lee Scholarship Fund at Seoul National University, Republic of Korea.

## Author Contributions

W-T Jeon, SSY Lee and Yuree Lee conceptualized the study; W-T Jeon, SSY Lee, C Sung, J-M Lee, A Cheon, B Kim, J-A Kim and Yeonu Lee carried out experiments; W-T Jeon, SSY Lee, C Sung, J-M Lee, A Cheon and B Kim conducted the investigation; W-T Jeon and A Cheon were responsible for data visualization; Yuree Lee, Y-S Noh and Y Joo acquired funding for the study; Yuree Lee, Y-S Noh, and Y Joo managed the project and provided supervision; W-T Jeon and Yuree Lee drafted the original manuscript; and all authors contributed to the review and editing of the manuscript.

## Data Availability

The RNA-Seq raw data have been deposited at the National Center for Biotechnology Information Sequence Read Archive under accession number PRJNA1505138.

## Supplemental Data

**Supplemental Table S1.** Normalized RNA-sequencing data from primary root tips after shoot excision

**Supplemental Table S2.** Gene Ontology enrichment analysis of gene clusters downregulated after shoot excision (FDR < 0.05; DAVID)

**Supplemental Table S3.** Expression rankings under intact conditions of genes downregulated after shoot excision

**Supplemental Table S4.** Expression patterns of intact-enriched, auxin-related, and cytokinin-related genes following shoot excision

**Supplemental Table S5.** Normalized RNA-sequencing data from LRS

**Supplemental Table S6.** Gene Ontology enrichment analysis of gene clusters upregulated in Zone 1

**Supplemental Table S7.** Gene Ontology enrichment analysis of gene clusters upregulated in Zone 1 following BA treatment

---

**Figure. S1.**
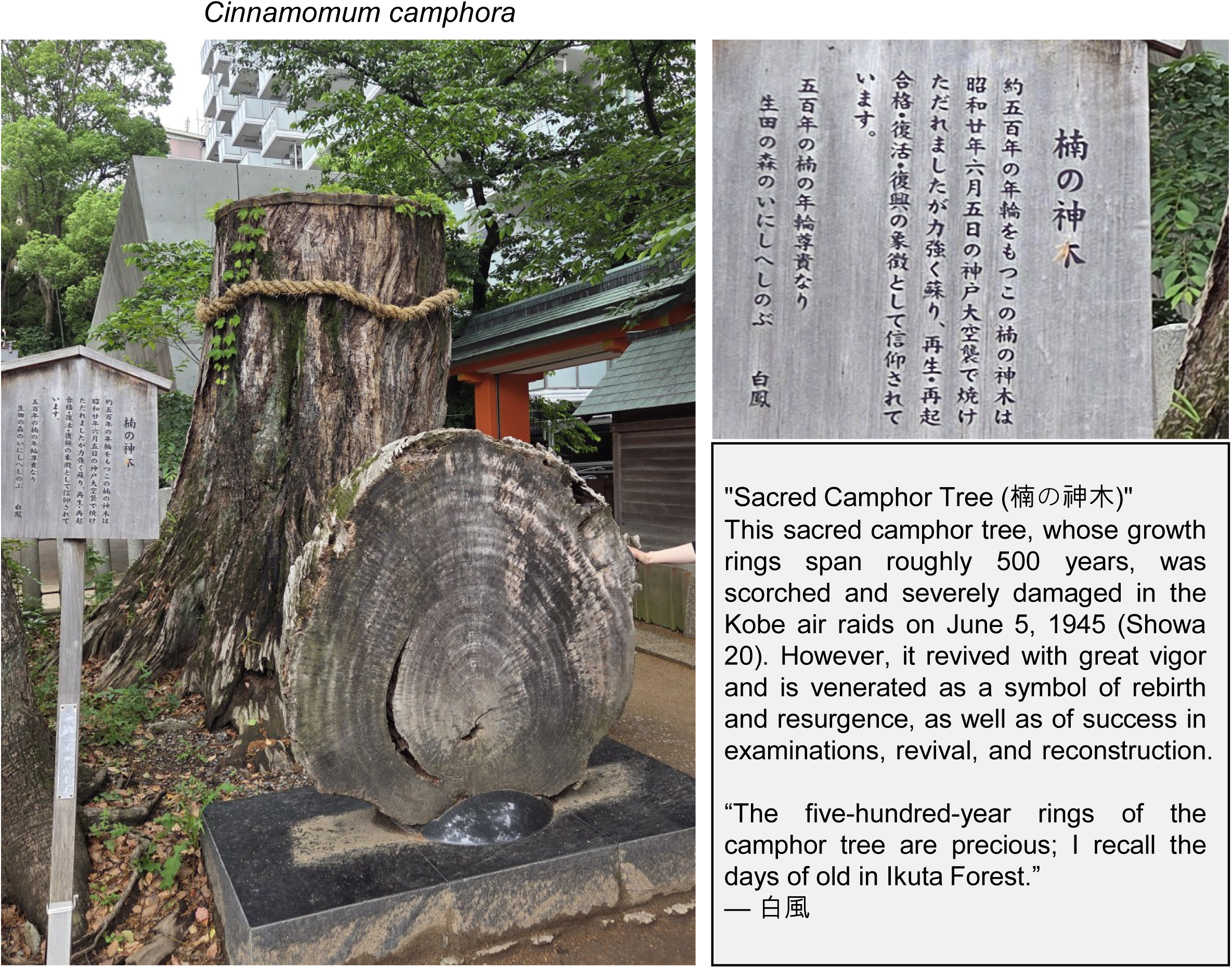
De-novo shoot regeneration after shoot excision. The sacred camphor tree, *Kusunoki no Shinboku*, at Ikuta Shrine in Kobe, Japan, together with the explanatory inscription describing the history of the tree. The inscription records that shoots have regenerated approximately 80 years after shoot excision.

**Figure. S2.**
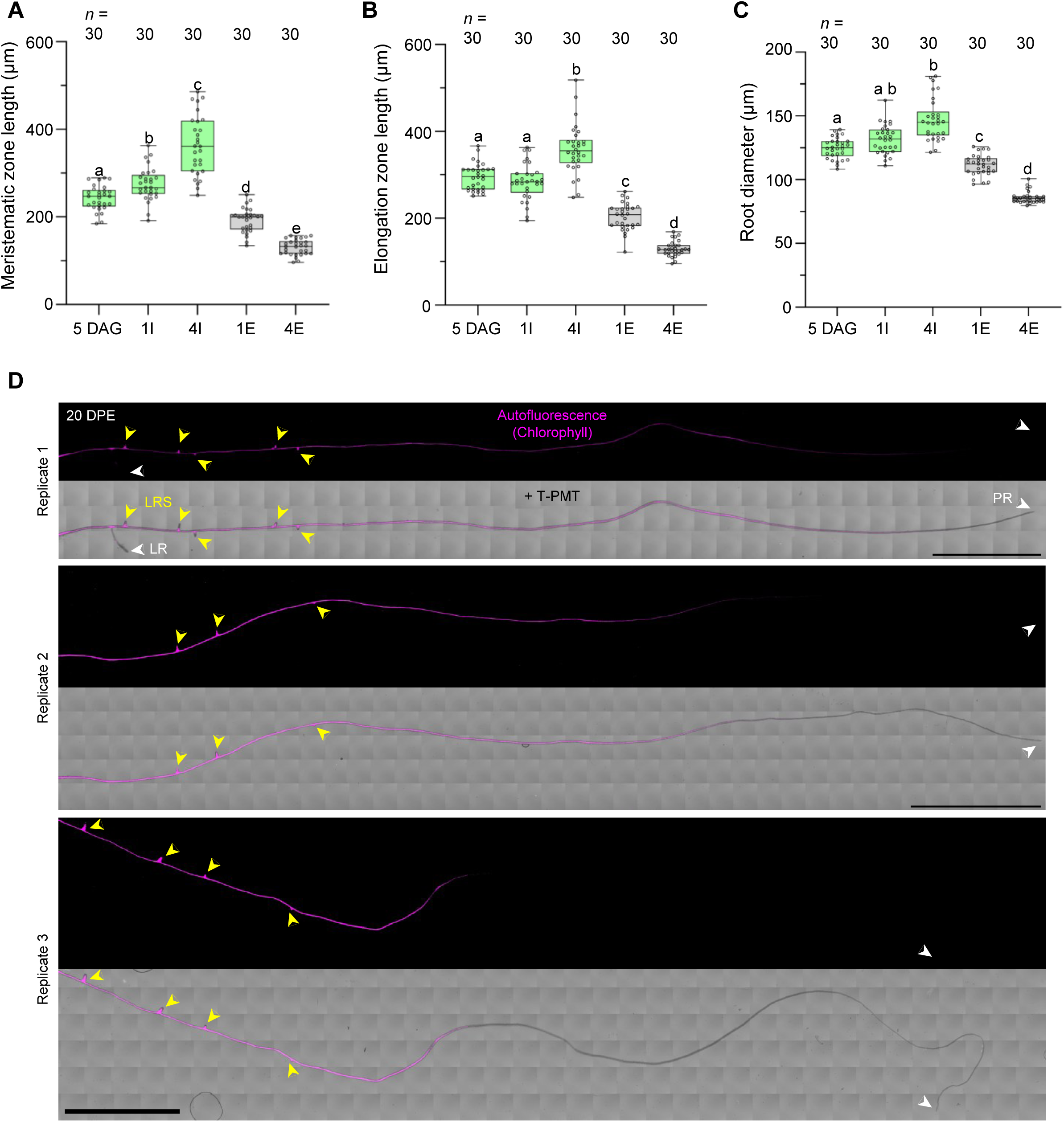
Remodeling of root architecture after shoot excision. (A to C) Changes in meristem zone length (A), elongation zone length (B), and root diameter (C) after shoot excision. I and E indicate intact and shoot-excised samples, respectively. Numbers preceding I or E indicate the number of days after 5 days after germination (DAG). Statistical significance was determined using Brown-Forsythe and Welch ANOVA tests followed by Dunnett’s T3 multiple comparisons test for (A) and (B), and the Kruskal-Wallis test followed by Dunn’s multiple comparisons test for (C). (D) Pattern of lateral root-derived shoot (LRS) formation after shoot excision. Yellow arrowheads indicate LRSs, and white arrowheads indicate lateral roots (LRs) and primary roots (PRs). DPE, days post excision. Scale bars, 5 mm.

**Figure. S3.**
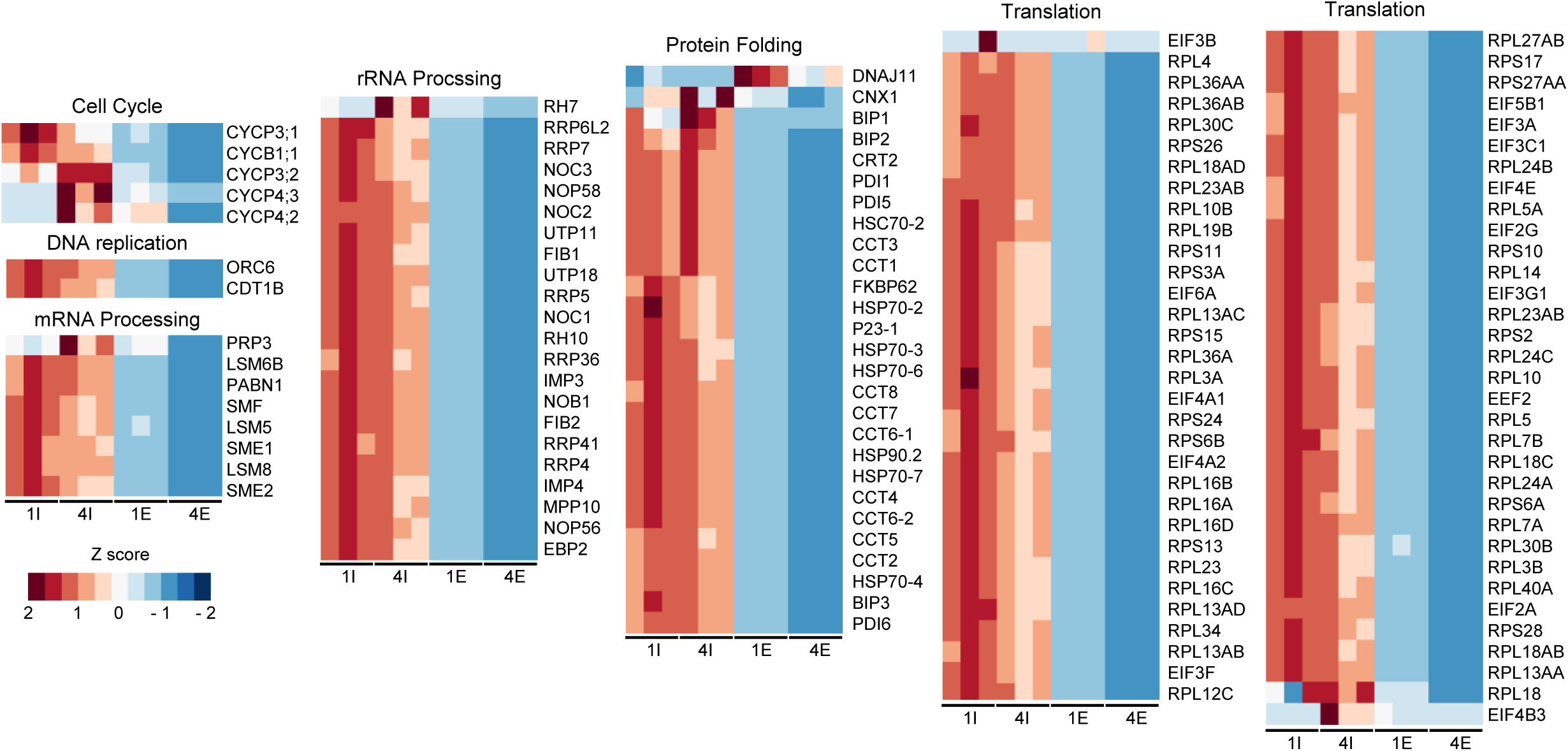
Transcriptional repression of cellular processes in root tips after shoot excision. Transcriptional repression of cellular processes in root tips after shoot excision. Heatmap of genes associated with Gene Ontology (GO) terms specifically downregulated in the excised group among differentially expressed genes (DEGs) identified in root tips (*P* < 0.05, fold change > 1.50). I and E indicate intact and shoot-excised samples, respectively.

**Figure. S4.**
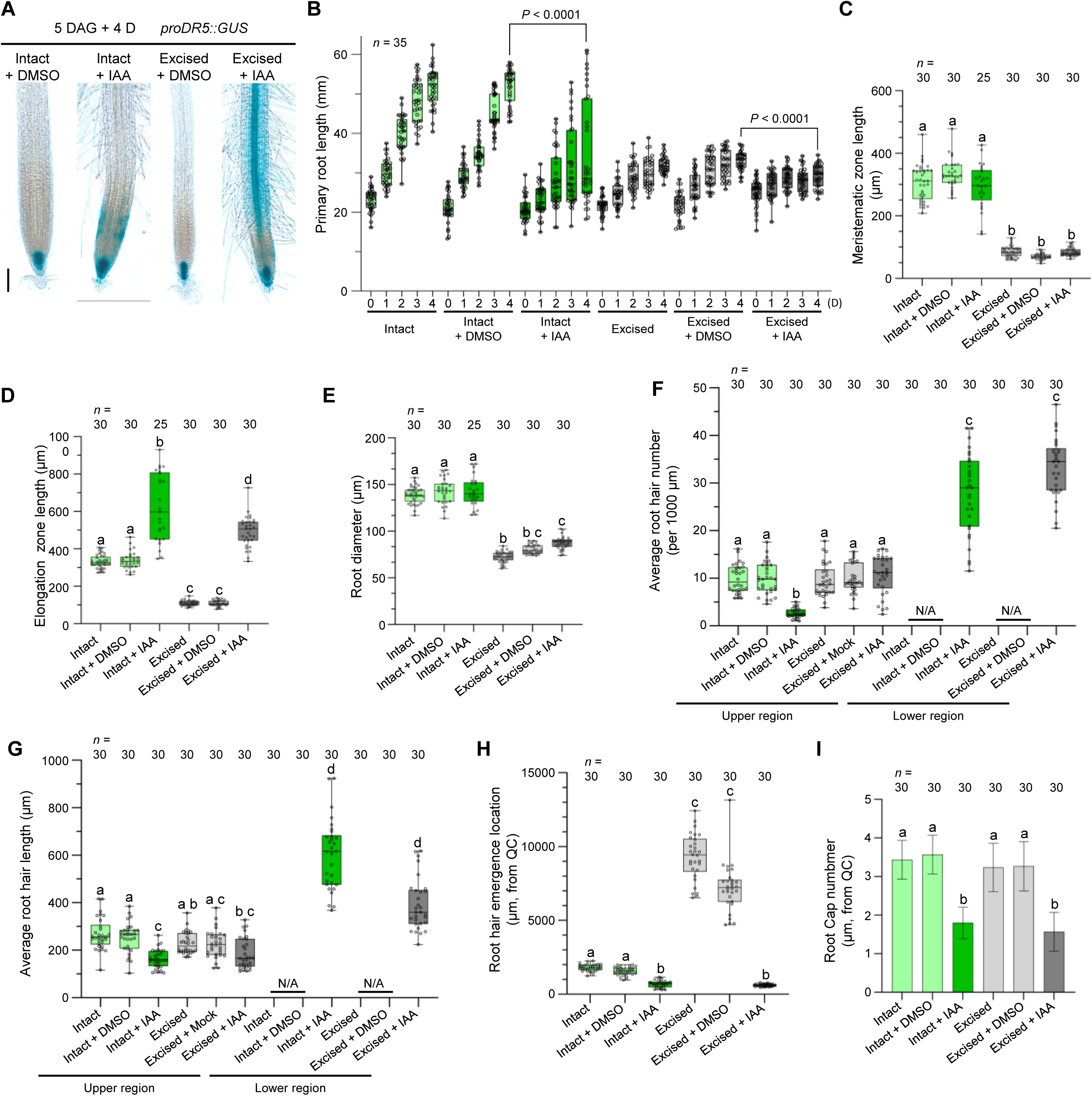
Responsiveness to exogenous IAA persists after shoot excision. (A) Representative images showing changes in *DR5::GUS* reporter activity in response to IAA treatment and shoot excision (*n* = 30). Scale bar, 100 μm. (B) Changes in primary root length in response to IAA treatment and shoot excision. (C to E) Changes in meristem zone length (C), elongation zone length (D), and root diameter (E) in response to IAA treatment and shoot excision. (F to H) Changes in root hair number (F), root hair length (G), and the root hair initiation site (H) in response to IAA treatment and shoot excision. The upper region was defined as the 5,000-μm region from 5,000 to 10,000 μm below the hypocotyl base. The lower region was defined as the 2,500-μm region from 0 to 2,500 μm above the quiescent center (QC). (I) Detached root cap number in response to IAA treatment and shoot excision. IAA treatment and shoot excision were performed at 5 days after germination (DAG). Statistical significance was determined using the Mann-Whitney test for the intact group in (B), Welch’s t test for the shoot-excised group in (B), and the Kruskal-Wallis test followed by Dunn’s multiple comparisons test for (E) to (I).

**Figure. S5.**
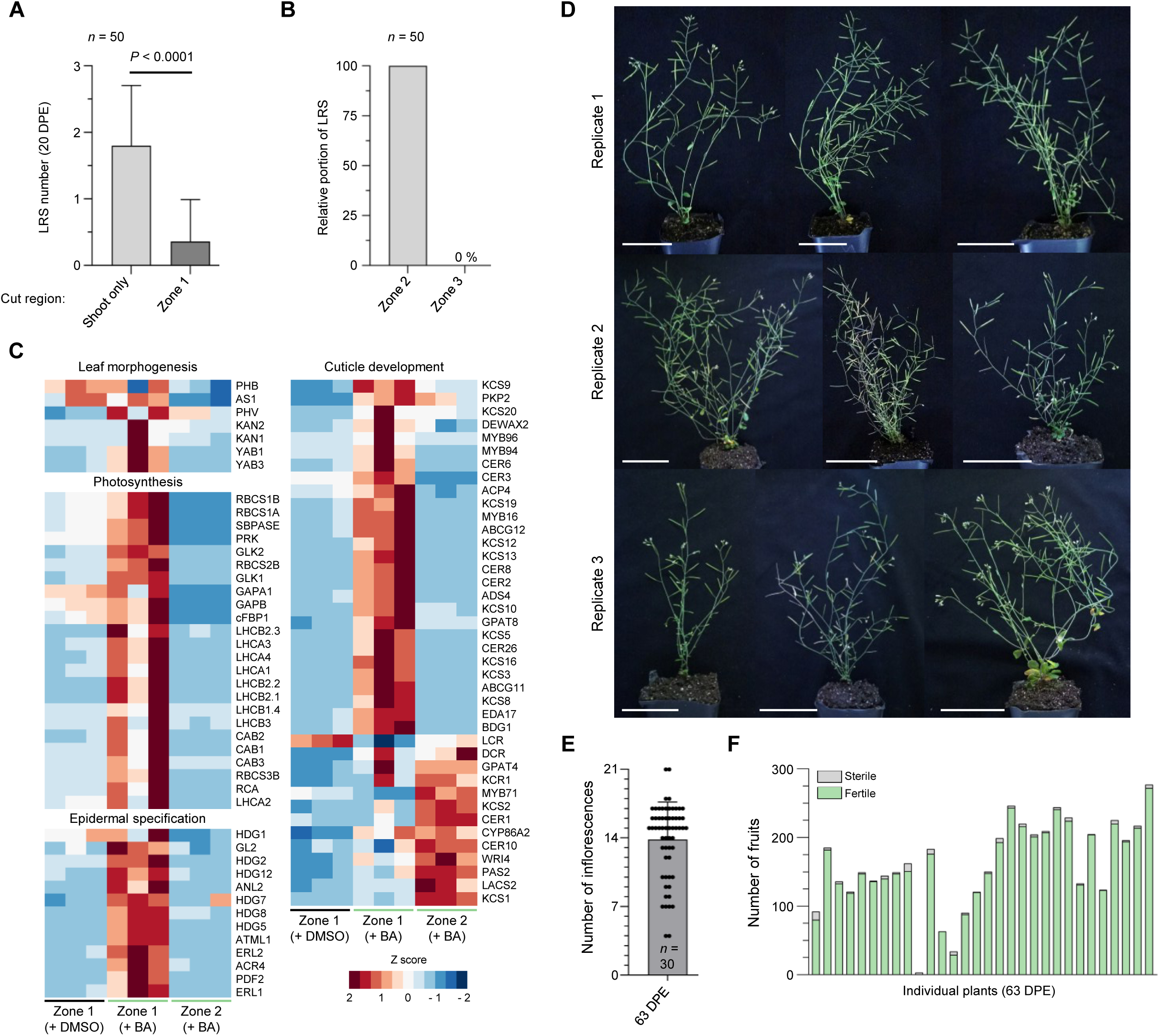
Exogenous cytokinin promotes shoot development in LRS. (A) Comparison of the number of LRSs formed after shoot excision alone or after excision of Zone 1, corresponding to the upper one-third of the primary root. Statistical significance was determined using the Mann-Whitney test. (B) Distribution of LRS formation sites after Zone 1 excision, showing whether LRSs formed in Zone 2, corresponding to the middle one-third of the primary root, or Zone 3, corresponding to the lower one-third of the primary root. (C) Heatmap showing the expression patterns of genes associated with leaf morphogenesis, photosynthesis, epidermal specification, and cuticle development among differentially expressed genes (DEGs; P < 0.05, fold change > 1.5). (D) Representative images of shoots developed at 69 days post excision (DPE) after shoot induction. Scale bars, 5 cm. (E and F) Quantification of inflorescence number (E) and fruit number (F) at 69 DPE. In (F), each bar represents an individual plant.

**Figure. S6.**
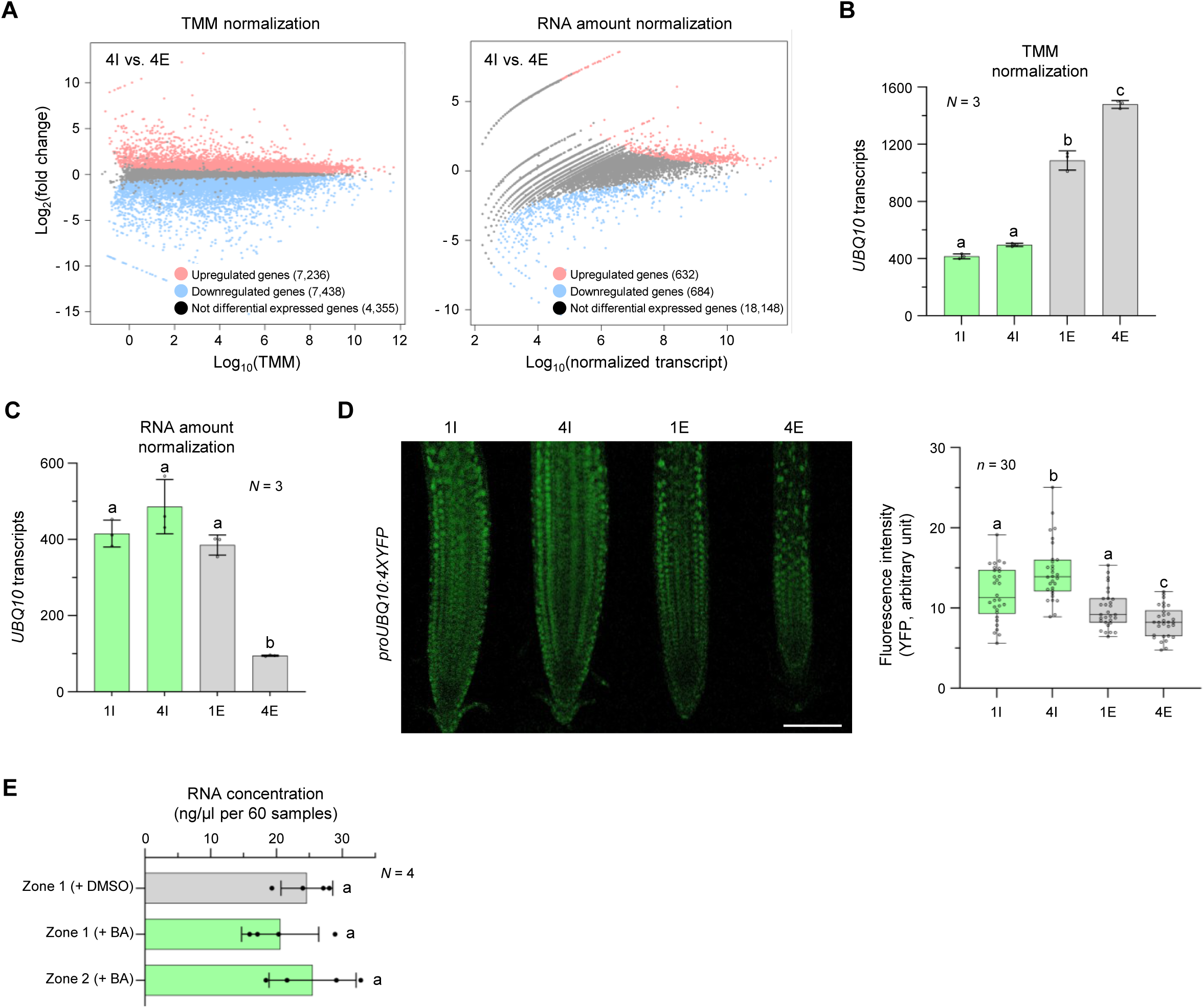
Normalization of root tip RNA-seq data after shoot excision. (A) Number of differentially expressed genes (DEGs) identified after normalization using either the trimmed mean of M values (TMM) method or RNA amount-based normalization. (B and C) Transcript abundance of UBQ10 in the RNA-seq data after TMM normalization (B) or RNA amount-based normalization (C). (D) Representative images and quantification of *proUBQ10:4xYFP* expression in shoot-intact and shoot-excised groups. Scale bar, 100 μm. (E) RNA concentration extracted from LRSs in different regions after treatment with DMSO or the exogenous cytokinin 6-benzylaminopurine (BA). Statistical significance was determined using ordinary one-way ANOVA for (B) and (E), and Brown-Forsythe and Welch ANOVA tests followed by Dunnett’s T3 multiple comparisons test for (C) and (D).

